# Trevolver: simulating non-reversible DNA sequence evolution in trinucleotide context on a bifurcating tree

**DOI:** 10.1101/672717

**Authors:** Chase W. Nelson, Yunxin Fu, Wen-Hsiung Li

## Abstract

**Summary:** Recent *de novo* mutation data allow the estimation of non-reversible mutation rates for trinucleotide sequence contexts. However, existing tools for simulating DNA sequence evolution are limited to time-reversible models or do not consider trinucleotide context-dependent rates. As this ability is critical to testing evolutionary scenarios under neutrality, we created Trevolver. Sequence evolution is simulated on a bifurcating tree using a 64 × 4 trinucleotide mutation model. Runtime is fast and results match theoretical expectation for CpG sites. Simulations with Trevolver will enable neutral hypotheses to be tested at within-species (polymorphism), between-species (divergence), within-host (*e.g*., viral evolution), and somatic (*e.g*., cancer) levels of evolutionary change.

**Availability and Implementation:** Trevolver is implemented in Perl and available on GitHub under GNU General Public License (GPL) version 3 at https://github.com/chasewnelson/trevolver.

**Supplementary information:** Further details and example data are available on GitHub.

## 1 Introduction

Coalescence simulation of gene trees provides a powerful, time-efficient approach for testing evolutionary hypotheses about observed patterns of genetic polymorphism and divergence. Programs exist for producing such trees under various models of neutrality, demographic change, migration, natural selection, recombination, and gene conversion (*e.g*., Hudson, 2002; Kelleher *et al*., 2016). After producing a tree, a common next step involves ‘seeding’ the root with an ancestral DNA sequence and simulating evolution under a given mutation model (Yang, 2014). Although excellent tools exist for this task (*e.g*., Cartwright, 2005), these programs are limited to standard DNA substitution models (*e.g*., General Time Reversible) or do not allow sequence context-dependent rates. Thus, it is currently not possible to use coalescence to test hypotheses that depend on trinucleotide context.

Sequence context strongly influences mutation (and thus substitution) rates at CpG sites, where methylation dramatically increases the CpG→TpG mutation rate in mammalian genomes (Hodgkinson and Eyre-Walker, 2011). As a result, derived CpGs have an elevated probability of back mutation, causing homoplasies (identity by state but not descent) that complicate evolutionary analysis. For example, homoplasies can lead to ancestral allele misclassification (polarization error) in the study of single nucleotide polymorphisms, causing artefactual patterns in distributions of derived allele frequencies (Hernandez et al. 2007). More generally, trinucleotide (3-mer; *i.e*., one flanking nucleotide on each side of a site) and other contexts have been shown to modulate mutation rates (Carlson *et al*., 2018). Because the rate of evolution expected under neutrality depends critically on mutational input, a tool is needed that accounts for context dependence when simulating evolution on a fixed tree, an approach that is more computationally efficient than forward-time simulation. We wrote Trevolver to fill this void.

## 2 Methods

Trevolver is written for Unix systems in Perl with no software dependencies. Input includes a bifurcating Newick tree, a seed sequence, a 64 × 4 trinucleotide mutation rate matrix, and the branch unit, a scaling factor used to obtain the number of generations on each branch.

The mutation rate matrix follows the format of SLiM 3 (Haller and Messer, 2019). Rows correspond to the 64 alphabetically-ordered trinucleotides (ancestral states) and columns correspond to the 4 alphabetically-ordered nucleotides (derived states of the central nucleotide). Each row of the mutation matrix should contain one value of 0 for the column corresponding to an identity, while the remaining values should be ≥0 and sum to the total mutation rate for the central nucleotide of that row. For example, the first row corresponds to AAA→AAA, AAA→ACA, AAA→AGA, and AAA→ATA rates. If the mean mutation rate for all sites is 0.9 × 10^-8^ per site per generation (*i.e*., no context-dependent variation), then the first column of the AAA row represents an identity and should be set to 0, while the remaining columns will each have the value 0.3 × 10^-8^, summing to 0.9 × 10^-8^. If the mutation model includes an elevated rate for CpGs, then rows containing a central CpG site (*e.g*., ACG) will sum to more than 0.9 × 10^-8^.

Evolution is simulated recursively. For each branch in the tree, the branch length is converted to generations, *g*. A random exponentially-distributed waiting time to the first mutation for the sequence is then calculated as *g*_w_ = -(1/*µ*) × ln(*x*_*1*_), where *µ* is the total mutation rate for the sequence, calculated by summing the context-dependent mutation rates for all sites, and *x*_*1*_ is a random number in the range [0,1]) (Yang, 2014). If the waiting time is less than the number of generations remaining on the branch (*i.e*., *g*_w_ < *g*), a mutation occurs. The site of the mutation is then chosen by generating a second random number (*x*_2_) in the range [0, *µ*]) and examining each trinucleotide in the sequence using a sliding window. If the current site has a mutation rate greater than 0, and *x*_*2*_ is less than the cumulative mutation rate of the sequence up to and including that site, a mutation occurs. Otherwise, the next site is examined. This process is continued until a site is found for which the above conditions hold, at which time a derived allele is selected with a probability equal to its relative rate. After the mutation, *µ* and *g* are recalculated to reflect the new trinucleotide content and time remaining on the branch. This allows back mutation to occur with a different (non-reversible) rate than forward mutation. Thus, the mutation rate can change over time and is an emergent property of the evolving sequence.

Trevolver internally stores the mutational history for every lineage and site. This information is inherited by descendant nodes, allowing the current sequence to be constructed at any time while maintaining space efficiency. Trevolver output includes the mutational history, a VCF file, and a multiple sequence alignment of the simulated sequences.

## 3 Results & Discussion

Trevolver was used to simulate the evolution of a 10,000 bp primate sequence for 10^7^ generations. The speed of execution was 137 mutations (16,421 generations) per second (2.9 GHz Intel Core i7). As expected, the number of CpGs increased dramatically under equal rates but decreased negligibly under more realistic, elevated-CpG rates, before reaching equilibrium (Fig. 1). Combined with coalescence simulations, Trevolver allows confidence intervals to be estimated for patterns of polymorphism expected under genetic drift of neutral mutations, enabling researchers to determine whether deviations from expectation are due to context-dependent mutation rate variation. Future developments could include wider odd-numbered *k*-mer contexts (*e.g*., 5-mer and 7-mer) and regional rate variation. Such developments should take advantage of the “one mutation, *k k*-mers” rule, *i.e*., one mutation will affect *k* existing polynucleotide strings of size *k* overlapping the site of change. This observation can be utilized to dramatically speed up computation, because an algorithm need only adjust the numbers of affected *k*-mers to compute the revised mutation rate for a sequence.

**Fig. 1.**
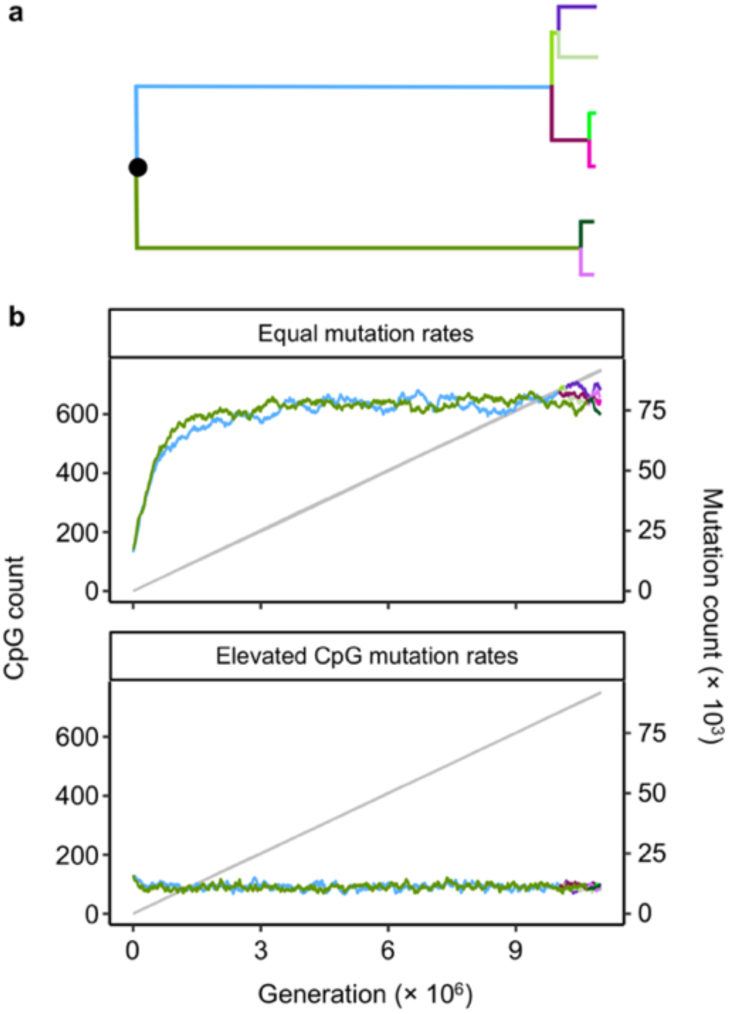
Simulated evolution of a 10,000-nucleotide primate sequence. Colors depict new lineages (branches) following a bifurcation (lineage split). Grey lines show number of mutations accumulated (right y-axis). (**a**) The fixed tree. (**b**) Equal mutation rates of 2.5 × 10^-7^ per site per generation (top) and context-dependent rates with a 20-fold increase to 5.0 × 10^-6^ at CpG sites (CGN and NCG trinucleotides) (bottom).

## Acknowledgments & Funding

This work was supported by a Gerstner Scholars Fellowship from the Gerstner Family Foundation at the American Museum of Natural History to C.W.N. We thank Reed A. Cartwright, Michael Dean, Dan Graur, Danielle A. Karyadi, Ming-Hsueh Lin, Lisa Mirabello, Michael Tessler, and Meredith Yeager for discussion.

